# Antibiotics promote susceptibility to *C. difficile* infection through a CCR5-dependent immune response

**DOI:** 10.64898/2026.06.24.734321

**Authors:** Md. Jashim Uddin, Nick R. Natale, Farha Naz, Jiayi Tian, Rachel McMillan, Duncan J. Hart, Shelby L. Schenck, William A. Petri

## Abstract

Antibiotics (ABXs) represent the current standard of care for treating *Clostridioides difficile* infection (CDI). Paradoxically, ABX-induced dysbiosis is the primary risk factor for CDI, as disruption of the colonic microbial ecosystem creates an opportunity for *C. difficile* colonization. Given that ABXs can also alter immune responses, we investigated whether ABXs prime the colonic immune milieu for CDI susceptibility. Here, we implicate ABXs in driving CDI severity through the emergence of pathogenic CCR5-reliant immune populations in the mouse colon. High-throughput immune cell profiling revealed that ABXs shift the colonic immune compartment toward a CCR5-associated type I immunity signature, marked by an expansion of CCR5^+^ ILC1s and CCR5^+^ Th1 cells. A partial genetic deletion of CCR5 reversed CDI severity, alleviating colonic inflammation and improving survival. Pharmacological inhibition of the CCL3/4/5-CCR5 circuit also recapitulated these favorable disease outcomes, which we attribute to reduced colonic CCR5^+^ ILC1, CCR5^+^ Th1, and CCR5^+^ CD8 T cell populations during CDI. Together, our findings extend beyond dysbiosis as the canonical CDI risk factor and establish ABX-induced immune imbalance as an underappreciated determinant of CDI susceptibility.

## INTRODUCTION

*Clostridioides difficile*, an opportunistic gram-positive, spore-forming anaerobe (1), continues to threaten global human health and strain contemporary healthcare systems. In the USA, hundreds of thousands of new cases of *C. difficile* infection (CDI) occur annually (2), resulting in burdensome costs for acute-care health facilities (3). If left untreated, infected individuals can succumb to severe colonic sepsis, with approximately 25,000 CDI-related deaths annually in the USA (4). Therefore, there is an urgent need to elucidate the pathomechanistic underpinnings of CDI to inform the development of new therapeutic strategies.

The resilience of *C. difficile* spores poses an alarming challenge as ingestion of disinfectant-resistant fecal-derived spores and subsequent germination in the colon results in symptoms ranging from mild diarrhea and abdominal pain to severe pseudomembranous colitis and toxic megacolon (5–8). The virulence factors of *C. difficile*, toxin A (TcdA) and toxin B (TcdB), glycosylate host Rho GTPases to inflict intense epithelial cell death and disrupt the intestinal barrier integrity (9, 10).

To date, there are few therapeutic options for treating CDI. The Infectious Diseases Society of America and Society for Healthcare Epidemiology of America (IDSA/SHEA) and the European Society of Clinical Microbiology and Infectious Diseases (ESCMID) recognize fidaxomicin or vancomycin as the preferred standard of care for first episodes of CDI (11). Oral metronidazole, formally considered the preferred standard of care, is recommended by the IDSA/SHEA and ESCMID only when fidaxomicin and vancomycin are unavailable (11). However, these ABX treatments fail to resolve the preexisting host conditions that initially permitted *C. difficile* to thrive.

Despite ABXs being regarded as the preferred standard of care for CDI, few risk factors are as consequential to CDI pathogenesis as excessive ABX usage. This phenomenon is primarily attributed to ABXs disrupting the native microbiota, thereby providing *C. difficile* with a competitive advantage for growth (7). In the laboratory, the standard CDI mouse model features a pre-treatment period of multiple ABXs, including gentamicin, colistin, metronidazole, vancomycin, and clindamycin (12). However, ABXs are also implicated in altering immune cell responses, independent of the microbiota (13, 14). For instance, the macrolide ABX family, which includes fidaxomicin, confers a wide range of immunomodulatory effects, including diminished pattern recognition receptor signaling, alterations in the cytokine secretome, and dampened antigen presentation (13, 14). Therefore, there is a critical need to further understand how ABXs shape host susceptibility.

In this study, we hypothesized that ABX-induced remodeling of the colonic immune landscape predisposes the host to severe CDI. Supporting this hypothesis, we found that ABX exposure led to the expansion of CCR5^+^ immune cell populations, specifically ILC1s, Th1 cells, and CD8 T cells. Abrogation of the CCL3/4/5-CCR5 circuit during CDI improved survival and attenuated inflammation, indicating that ABXs provoke a pathogenic immune imbalance in the colon through CCR5 signaling. Overall, this study sheds new light on the immunological consequences of excessive ABX use and brings ABXs as the standard of care for CDI under renewed scrutiny.

## RESULTS

To assess whether ABX exposure alters the colonic immune compartment, we administered to naïve mice either a sham treatment regimen or a commonly used ABX regimen that serves as a standard pre-treatment in CDI mouse models (12). Mouse colons were harvested three days after clindamycin injection and underwent flow cytometry analyses to capture the state of the colonic immune compartment that exists before a hypothetical *C. difficile* challenge (Fig. 1a and S1). In the colons of ABX-treated mice, we detected a marked expansion of CD45^+^ cells (Fig. S1; Fig. S2a). While myeloid populations remained unchanged (Fig. S2b-f), ILC1s, Th1 cells, and CD8 T cells were elevated in the colon of ABX-treated mice (Fig. 1b-d). In particular, ABX exposure was associated with an increased abundance of colonic CCR5^+^ ILC1s (Fig. 1e) — an ABX-inducible response that we have previously characterized (15). CCR5^+^ Th1 cells and CCR5^+^ CD8 T cells, albeit to a lesser extent, were also expanded (Fig. 1f and 1g), which suggests that ABX treatment establishes a type I immune profile in the colon. Whether this ABX-induced type I immune profile is a driver of CDI susceptibility became a major focus of our research.

**Fig. 1.**
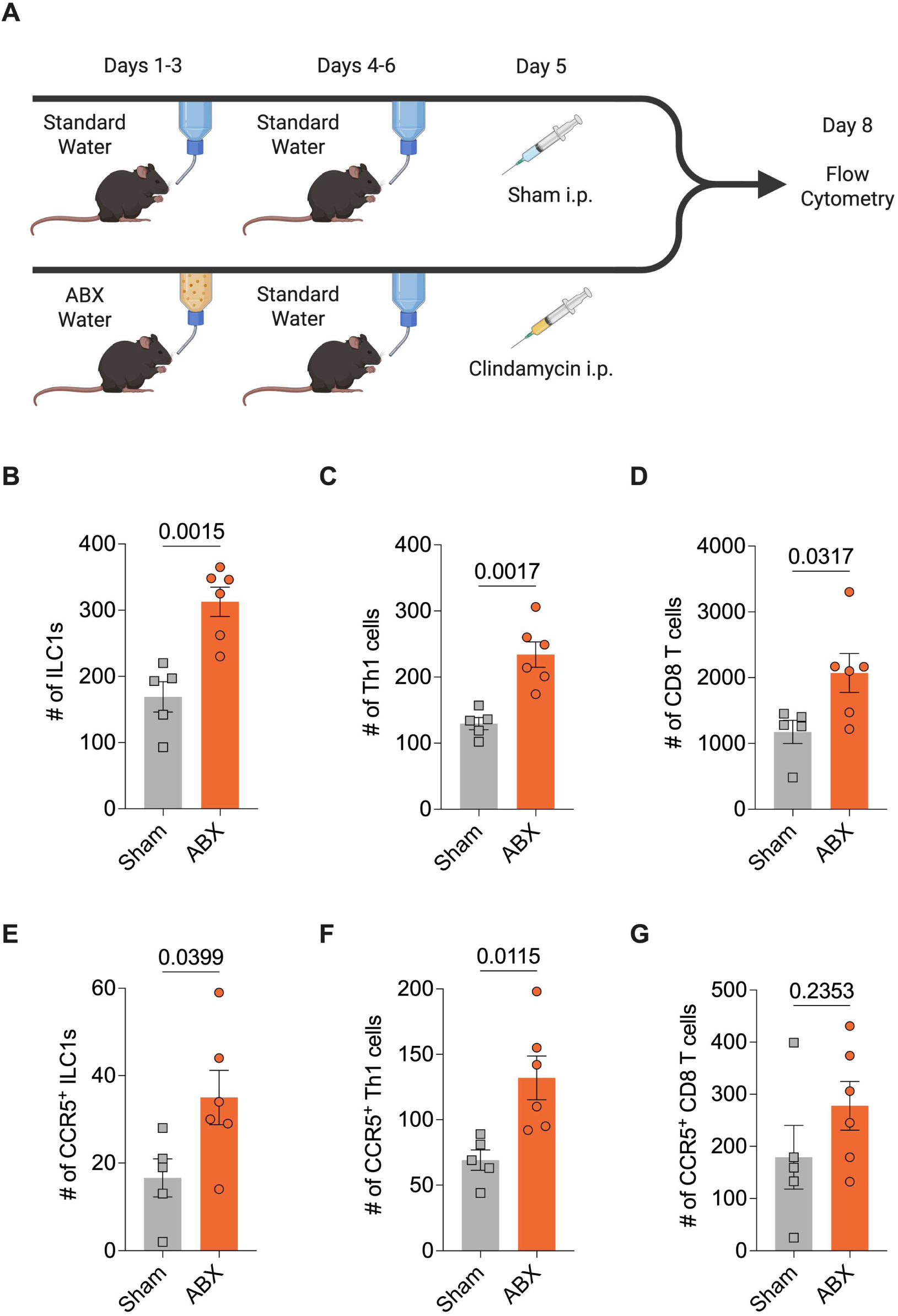
ABXs promote the expansion of a CCR5-associated type I immune profile in the colon. (**A**) Schematic diagram of experimental design. C57BL/6J mice received (1) either normal drinking water (sham treatment) or drinking water supplemented with an ABX cocktail of metronidazole, vancomycin, gentamicin, and colistin (ABX treatment) and (2) an i.p. injection of either a sham solution (sham treatment) or clindamycin (ABX treatment). The colon of each mouse was harvested 7 days after the start of the ABX regimen. (**B-G**) Cell count quantification of ILC1s (B), Th1 cells (C), CD8 T cells (D), CCR5^+^ ILC1s (E), CCR5^+^ Th1 cells (F), and CCR5^+^ CD8 T cells (G) per harvested colon. Statistics calculated by a Welch’s t-test. Data are presented as mean (± SEM). Each symbol represents a value from an individual animal.

Since many of the expanded type I immunity-associated cells in the ABX-treated colon expressed CCR5, we reasoned that the recruitment of peripheral immune cells to the colon via a CCL3/4/5-CCR5 chemotactic signaling pathway could be a driver of CDI immunopathology. To test this, wild-type (WT) mice and heterozygous CCR5 knockout mice (CCR5^+/-^) were challenged with *C. difficile*. Despite CCR5^+/-^ mice exhibiting no rescue in weight loss at early disease timepoints (Fig. 2a), these mice had lower clinical scores at early disease timepoints and a lower probability of succumbing to CDI (Fig. 2b and c). In addition, CCR5^+/-^ mice had longer colons, a macroscopic indicator of reduced inflammation (16), compared to wild-type controls (Fig. 2d and e).

**Fig. 2.**
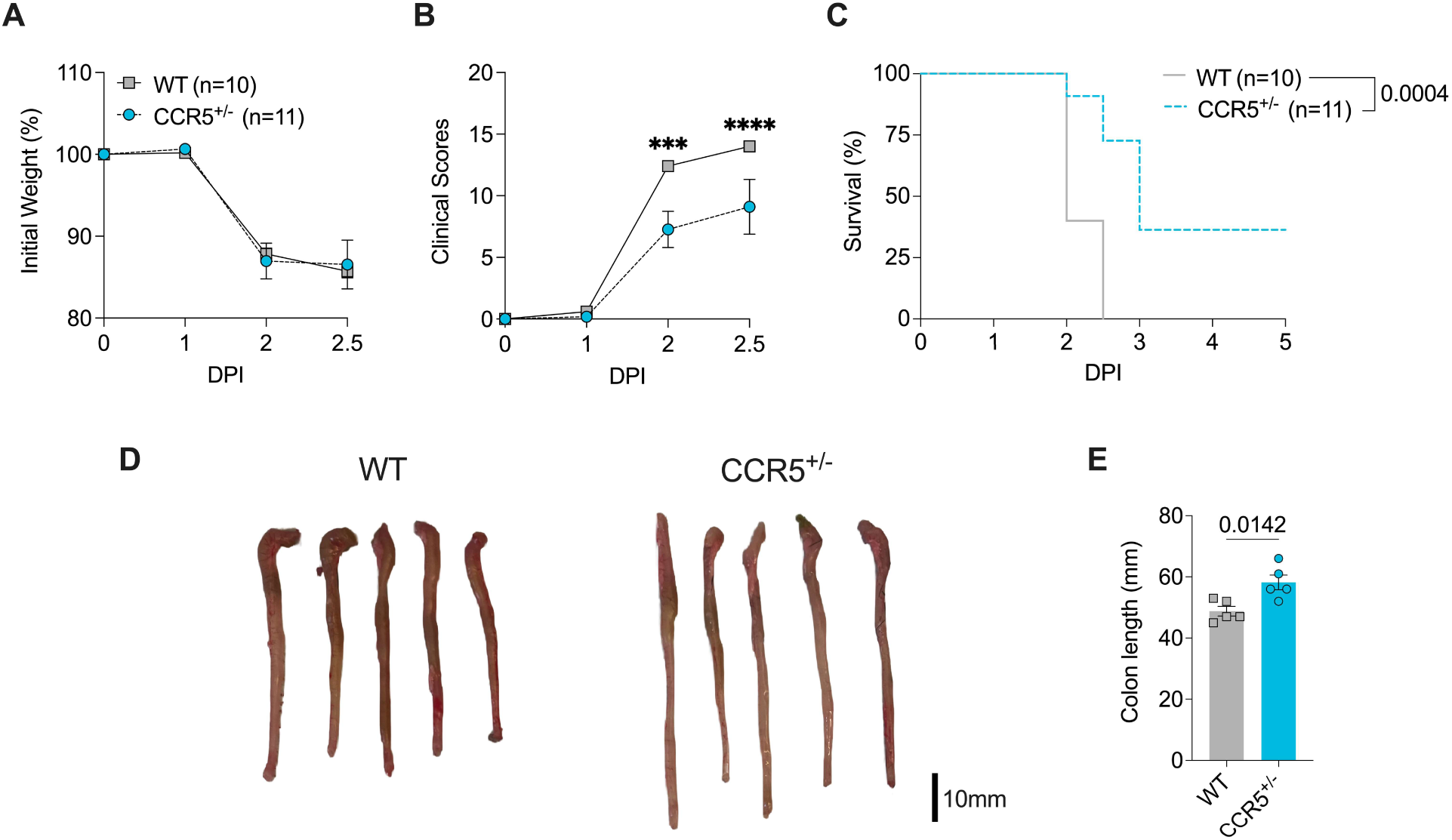
A partial genetic deletion of CCR5 dampens CDI severity. WT (C57BL/6J) and CCR5^+/-^ mice received drinking water supplemented with an ABX cocktail of metronidazole, vancomycin, gentamicin, and colistin and an i.p. injection of clindamycin. Mice were infected with 1,000 *C. difficile* spores via oral gavage 24h after i.p. injection of clindamycin. (**A**) Change in body weight over time. (**B**) Clinical scores over time. (**C**) Kaplan-Meier curve displaying the percentage of mice surviving over time. (**D**) Representative mouse colons. (**E**) Quantification of colon length on DPI 2. Each symbol represents a value from an individual animal. (A, B, C) Pooled data from two independent experiments. (A, B, C, E) Statistics calculated by linear GEE with Holm’s multiple testing correction (A), a negative binomial GEE with Holm’s multiple testing correction (B), a Log-Rank test (C), and a Welch’s t-test (E). Statistics calculated by a Welch’s t-test. Data are presented as mean (± SEM). ***P < 0.001, ****P < 0.0001.

To further interrogate the role of the CCL3/4/5-CCR5 axis in CDI pathogenesis, we employed maraviroc, a CCR5 inhibitor that binds to an allosteric pocket of CCR5 to prevent CCL3/4/5-directed chemotaxis (17, 18). *C. difficile*-challenged C57BL/6J mice received daily i.p. administration of either the vehicle or maraviroc (Fig. 3a). Consistent with the phenotypic outcomes observed with CCR5^+/-^ mice, maraviroc-treated mice presented with no difference in weight loss trajectories at early disease timepoints but exhibited lower clinical scores and improved survival (Fig. 3b-f). These findings build on existing evidence that blocking CCR5 protects against colitis (19) and suggest that maraviroc may have therapeutic potential for treating CDI.

**Fig. 3.**
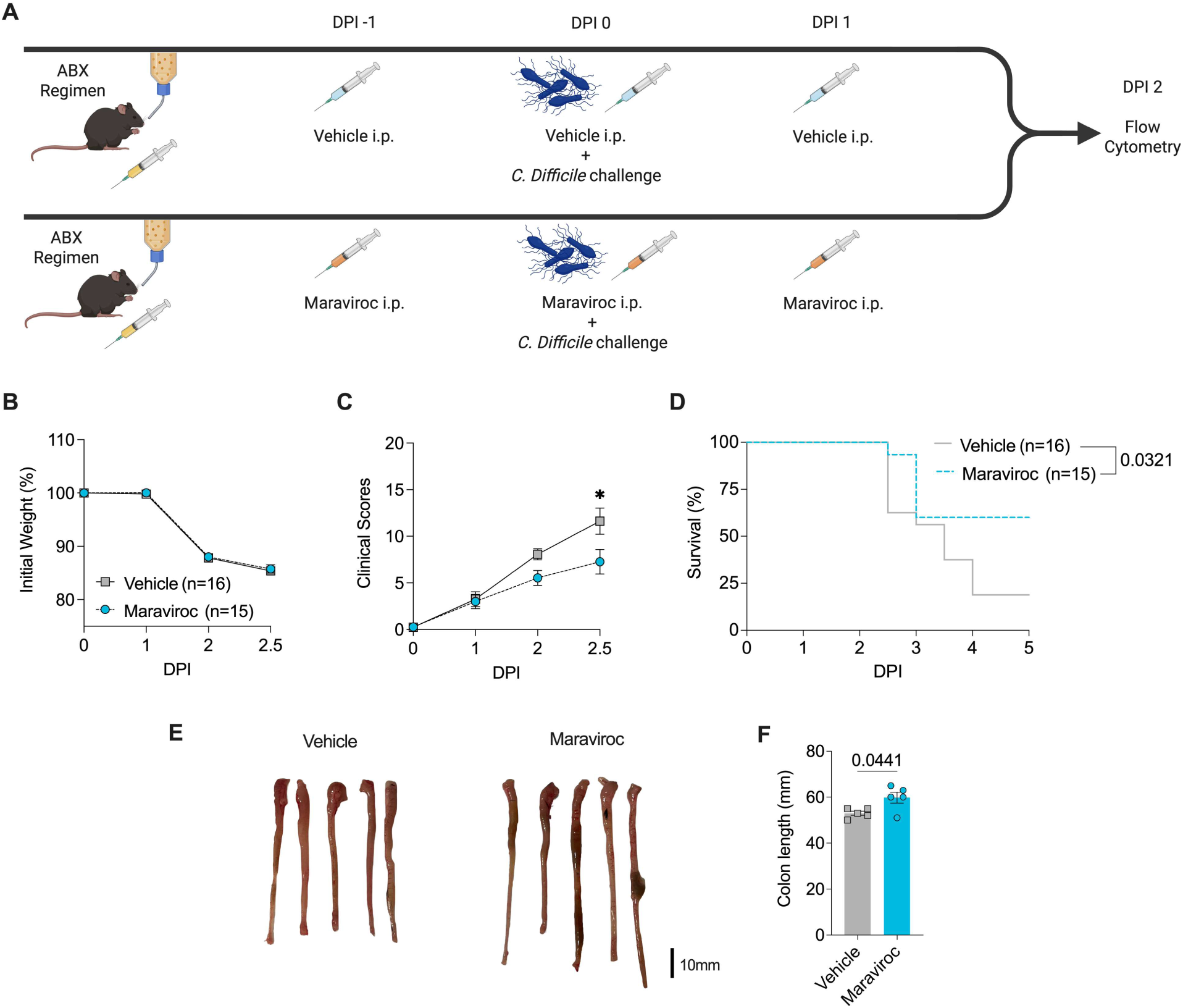
Therapeutic CCR5 inhibition mitigates CDI severity. (**A**) Schematic diagram of experimental design. C57BL/6J mice received drinking water supplemented with an ABX cocktail of metronidazole, vancomycin, gentamicin, and colistin and an i.p. injection of clindamycin. Mice were infected with 1,000 *C. difficile* spores via oral gavage 24h after i.p. injection of clindamycin. On DPI-1, DPI 0, and DPI 1, mice were treated via i.p injections with either 1x PBS (vehicle) or a CCR5 inhibitor (maraviroc). (**A**) Change in body weight over time. (**B**) Clinical scores over time. (**C**) Kaplan-Meier curve displaying the percentage of mice surviving over time. (**E**) Representative mouse colons. (**F**) Quantification of colon length on DPI 2. Each symbol represents a value from an individual animal. (B, C, D) Pooled data from two independent experiments. (B, C, D, F) Statistics calculated by a linear GEE with Holm’s multiple testing correction (B), a negative binomial GEE with Holm’s multiple testing correction (C), a Log-Rank test (D), and a Welch’s t-test (F). Data are presented as mean (± SEM). *P < 0.05.

Given that ABXs promote a CCR5-associated type I immunity signature in the colon and that abrogation of the CCL3/4/5-CCR5 axis yields favorable CDI outcomes, we sought to examine whether CDI rescue was accompanied by an attenuation in ABX-influenced type I immune populations. Flow cytometry analyses revealed that the number of colonic CCR5^+^ ILC1s, CCR5^+^ Th1 cells, and CCR5^+^ CD8 T cells decreased in maraviroc-treated mice – a colonic immune profile that is strikingly opposite to ABX exposure (Fig. 4a-f). This downturn in CCR5^+^ ILC1, CCR5^+^ Th1, CCR5^+^ CD8 T cell prevalence, which coincided with improved CDI outcomes, is concordant with our hypothesis that blunting the CCR5-dependent type I immune response results in protective outcomes during CDI. Conversely, maraviroc-treated mice had an expanded population of eosinophils (Fig. S3a and S3b), a cell population previously linked to protective CDI outcomes (20, 21). Together, these data suggest that CCR5 inhibition rebalances the colonic immune compartment away from type I immunity and toward a potentially protective eosinophilic response.

**Fig. 4.**
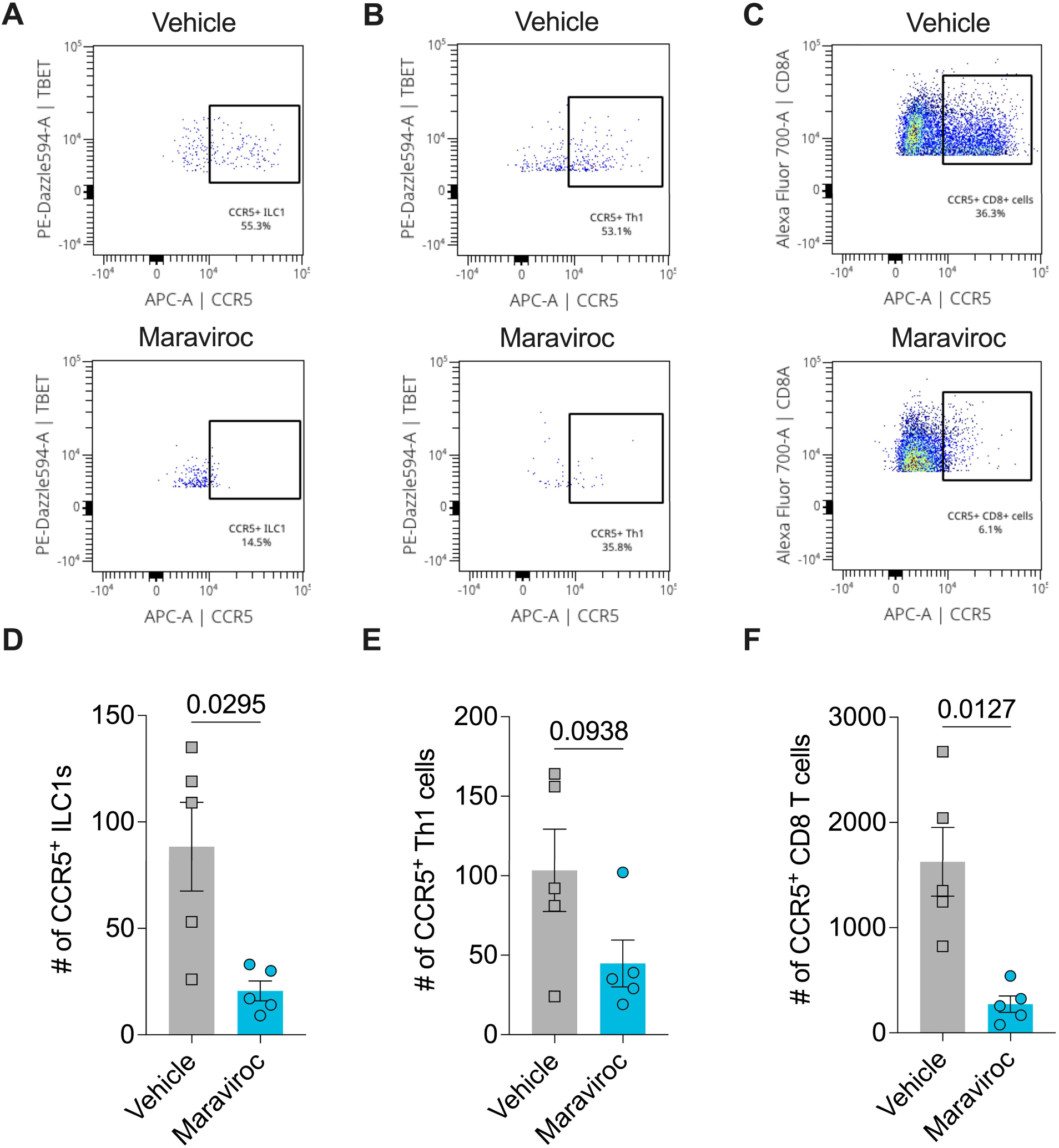
Therapeutic CCR5 inhibition curbs a CCR5-associated type I immune profile in the colon during CDI. C57BL/6J mice received drinking water supplemented with an ABX cocktail of metronidazole, vancomycin, gentamicin, and colistin and an i.p. injection of clindamycin. Mice were infected with 1,000 *C. difficile* spores via oral gavage 24h after i.p. injection of clindamycin. On DPI-1, DPI 0, and DPI 1, mice were treated via i.p injections with either 1x PBS (vehicle) or a CCR5 inhibitor (maraviroc). The colon of each mouse was harvested on DPI 2 (7 days after the start of the ABX regimen). (**A-C**) Representative flow cytometry dot plots showing the gating strategies for CCR5^+^ ILC1s (A), CCR5^+^ Th1 cells (B), and CCR5^+^ CD8 T cells (C). (**D-F**) Cell count quantification of CCR5^+^ ILC1s (D), CCR5^+^ Th1s (E), and CCR5^+^ CD8 T cells (F) per harvested colon. Statistics calculated by a Welch’s t-test. Data are presented as mean (± SEM). Each symbol represents a value from an individual animal.

## DISCUSSION

In this study, we provide *in vivo* evidence that ABX exposure reshapes the colonic immune landscape — a major consequence that contributes to CDI susceptibility. ABX treatment enriched the colon with type I immunity-associated leukocytes, including CCR5^+^ ILC1s and CCR5^+^ Th1 cells, prompting us to investigate the role of CCR5 in orchestrating the immune response during CDI. Both partial genetic deletion of CCR5 and pharmacological inhibition of CCR5 ameliorated disease severity, suggesting that a CCR5-dependent immune circuit drives CDI susceptibility. Collectively, these findings unveil a CCR5-reliant type I immune response as an underappreciated mechanism by which ABXs prime the host for severe CDI.

Despite establishing a causative link between CCR5-dependent type I immunity and CDI severity, the mechanistic drivers underlying this ABX-induced immune imbalance remain unclear. The loss of microbial-immune crosstalk following ABX treatment may contribute to this phenomenon, as the microbiota maintains immune homeostasis (22–24) and facilitates dynamic shifts in T-helper cell subsets (25–28). Consistent with observations by other research groups (27, 28), disruption of the homeostatic microbiota with ABXs biased the CD4^+^ T cell compartment toward a Th1 profile in our model prior to enteric infection. This expansion of the Th1 population following ABX treatment could be due to the propensity of ABX to dysregulate antigen-presenting cells, which has been observed *in vitro* (29–31) and *in vivo* (27, 28). However, we also detected an enrichment of ILC1s following ABX treatment, which do not rely on antigen presentation for activation, thus challenging our assumption that the expansion of the type I immune profile was the result of antigen-presenting cell dysregulation. ABX-induced remodeling of the chemokine secretome of colonic antigen-presenting cells may promote the coordinated infiltration of both CCR5^+^ ILC1s and CCR5^+^ Th1 cells into the tissue. Whether ABX-induced dysbiosis fosters a colonic niche enriched in chemotactic ligands for CCR5 remains an important question for future investigation.

The purpose of CCR5^+^ immune cell expansion in the colon during dysbiosis remains unknown. Since ABX use depletes commensal microbial populations in the colon, it creates an opportunity for ABX-resistant pathogenic bacteria, normally constrained by commensal microbes by competitive exclusion, to proliferate (7). We propose that the expansion of a CCR5^+^ immune compartment in the colon constitutes a compensatory host response to heighten immunosurveillance of impending colonization by opportunistic pathogenic bacteria. The type I immunity–biased profile of colonic immune cells following ABX treatment suggests a particular focus on protecting against intracellular bacterial pathogens, such as *Listeria*, *Salmonella*, and *Shigella*. Since *C. difficile* is an extracellular bacterium that is primarily curbed through the host’s type III immunity processes, it is unsurprising that the expansion of type I immunity-related cell populations exacerbated CDI severity.

In sum, the findings of this study indicate that the immune axes underlying CDI pathogenesis remain largely unexplored. Although the CDI research community has made great strides in developing anti-TcdA and anti-TcdB monoclonal antibodies to confer robust CDI protection (32, 33), the potential emergence of new ABX-resistant, hypervirulent *C. difficile* strains demands alternative therapeutic approaches (34). Therefore, the CCR5-dependent type I immune response during CDI warrants further investigation: a deeper mechanistic understanding of this immunopathogenic profile could uncover targetable pathways for future immunotherapies designed to mitigate CDI severity in the clinic.

## METHODS

### Mice

All mice were male and underwent experimentation at 8-12 weeks of age. Wild-type (WT; C57BL/6J) mice were purchased from The Jackson Laboratory. CCR5^+/-^ mice were generated by breeding C57BL/6J mice with CCR5^-/-^ mice at the University of Virginia vivarium. To equilibrate the microbiomes, bedding was shared between WT mice and CCR5^+/-^ mice for at least 2 weeks before the start of the experiments. Mice were kept under specific-pathogen-free conditions and provided autoclaved food and water. All experimental procedures were approved by the University of Virginia Institutional Animal Care and Use Committee.

### Antibiotic treatment

To make mice susceptible to *C. difficile* infection, all mice were given antibiotics starting 6 days before the *C. difficile* challenge. Briefly, an antibiotic cocktail (45 mg/L vancomycin, 35 mg/L colistin, 35 mg/L gentamicin, and 215 mg/L metronidazole) was added to the drinking water and continued for 3 days. After 3 days, antibiotic water was replaced with regular drinking water. Two days later (1 day before the *C. difficile* challenge), mice received an i.p. injection of clindamycin (0.016 mg/g; Hospira) diluted in saline. The sham group was supplied with regular drinking water and received a concomitant i.p. saline injection administered alongside the ABX-treated mice.

### C. difficile challenge

Antibiotic-treated mice were infected with 1,000 *C. difficile* (R20291 strain) spores diluted in 100 µL sterile water via oral gavage.

### Clinical scoring

Mice were observed daily for disease signs, and clinical scores were recorded. Clinical scores were determined by assessing weight loss, coat appearance, posture, eye condition, activity, and diarrhea. For weight loss and activity, mice were scored from 0 to 4. For coat, posture, eyes, and diarrhea, scores ranged from 0 to 3 for each parameter. Mice were considered to have reached euthanasia criteria and were humanely euthanized if weight loss and activity scores reached 4 or if the total combined score was ≥14. If a mouse was found dead in the cage, a total clinical score of 20 was assigned.

### Maraviroc treatment

To prepare the CCR5 inhibitor solution, maraviroc (SelleckChem #UK-427857) was first dissolved in DMSO (Final concentration: 5% DMSO) and subsequently diluted in 1x PBS (vehicle). Each mouse received 1 mg of maraviroc via i.p. injections on DPI −1, 0, and 1 of relative to *C. difficile* challenge.

### Preparation of single-cell suspension

Mice were harvested in accordance with an approved ACUC protocol, and colons were collected and measured in length with a standard ruler. A longitudinal excision was made along the colon to remove its contents. Cleaned tissues were washed in buffer 1 (HBSS with 25 mM HEPES and 5% FBS) and stored in buffer 1 until all mice were harvested. Tissues were then incubated in buffer 2 (HBSS with 15 mM HEPES, 5 mM EDTA, 10% FBS, and 1 mM DTT) for 40 min at 37 °C with shaking at 220 rpm to remove the epithelial layer from the lamina propria. After that, the lamina propria was collected and dissected into small pieces. Dissected tissues were incubated in digestion buffer (RPMI 1640 containing 0.17 mg/mL Liberase TL and 30 µg/mL DNase) for 30 min at 37 °C with shaking at 220 rpm. The digested tissues were then passed through 100 µm and 40 µm cell strainers. Cells were pelleted by centrifugation at 500 × g for 5 min and reconstituted in FACS buffer (2% FBS in PBS).

### Flow cytometry

For flow cytometry staining, cells were plated in a 96-well round-bottom plate. Cells were first stained for live/dead discrimination by incubating with 1 µL of Zombie NIR (BioLegend, 423105) in 100 µL of PBS for 30 minutes on ice or in a refrigerator. Excess reagent was removed by washing twice with FACS buffer. Cells were then surface-stained with antibodies against LY6C (Alexa Fluor 488, BioLegend, 128022), LY6G (BV650, BioLegend, 127641), Siglec-F (PE, BioLegend, 155506), CD45 (Spark Violet 538, BioLegend, 103180), CD8a (AF700, BioLegend, 100730), CD4 (APC-Fire 750, BioLegend, 100460), CD11c (PE-Cy7, BioLegend, 117318), CD11b (BV480, Fisher Scientific, 566117), CD3 (APC, BioLegend, 100235), TCRβ (BV570, BioLegend, 109231), CD90.2 (BV785, BioLegend, 105331), CD127 (PE-Cy5, BioLegend, 135016), CXCR3 (BV605, BioLegend, 126523), CD64 (BV421, BioLegend, 139309), and CCR5 (APC, BioLegend, 107012). For intracellular staining, cells were fixed and permeabilized using the FoxP3/Transcription Factor Staining Buffer Set (eBioscience, 00-5523-00) according to the manufacturer protocol. Cells were then stained with antibodies against T-bet (PE/Dazzle 594, BioLegend, 644828), GATA3 (BV711, BD, 565449), and RORγT (PerCP eFluor 710, eBioscience, 46-6981-82). Samples were run at the University of Virginia Flow Cytometry Core using a 5-laser Cytek Aurora Borealis flow cytometer.

## Statistical analysis

Changes in body weight, which were continuous and approximately Gaussian in distribution, were analyzed using linear generalized estimating equations (GEEs) with Holm’s multiple testing correction to compare differences between groups across repeated measurements. Clinical scores were analyzed with negative binomial GEEs with Holm’s multiple testing correction. All GEEs were modeled with R (v.4.3.1) using RStudio (Posit, v.2023.06.1-524) software. For analyses of animal survival, a log-rank (Mantel–Cox) test was performed in GraphPad Prism (v.10.1.1) software. Welch’s t-tests were conducted for all flow cytometry and colon length analyses in GraphPad Prism (v.10.1.1) software. P values lower than 0.05 were considered significant. *P < 0.05, ***P < 0.001, ****P < 0.0001.

## ACKNOWLEDGEMENTS

This work was supported by grants from the National Institutes of Health (NIH) that were awarded to William A. Petri, Jr. (R01AI124214, R01AI152477, 5R37AI026649). Nick R. Natale was supported by the Global Biothreats Training Grant (NIH T32AI055432), the UVA Brain Institute Presidential Fellowship in Collaborative Neuroscience, and the Ruth L. Kirschstein National Research Service Award (NRSA) Individual Predoctoral Fellowship (NIH F31NS135897). Rachel McMillan was supported by the Global Biothreats Training Grant (NIH T32AI055432). We thank the UVA Flow Cytometry Core (RRID: SCR_017829) for their technical support during the data collection process. BioRender.com was used to generate all figure schematics. GraphPad Prism (v.10.1.1) was used to visualize all data.

## AUTHOR CONTRIBUTIONS

Md. Jashim Uddin, Conceptualization, Data curation, Formal analysis, Investigation, Validation, Visualization, Writing – original draft, Writing – review and editing | Nick R. Natale, Conceptualization, Data curation, Formal analysis, Investigation, Visualization, Writing – original draft, Writing – review and editing | Farha Naz, Investigation, Writing – review and editing | Jiayi Tian, Investigation, Writing – review and editing | Rachel H. Boone, Investigation, Writing – review and editing | Duncan J. Hart, Investigation, Writing – review and editing | Shelby L. Schenck – Investigation, Writing – review and editing | William A. Petri Jr., Conceptualization, Funding acquisition, Project administration, Supervision, Writing – review and editing

## DATA AVAILABILITY

Any requests for raw data can be directed to Md. Jashim Uddin and William A. Petri, Jr.

---

**Supplementary Fig. 1.**
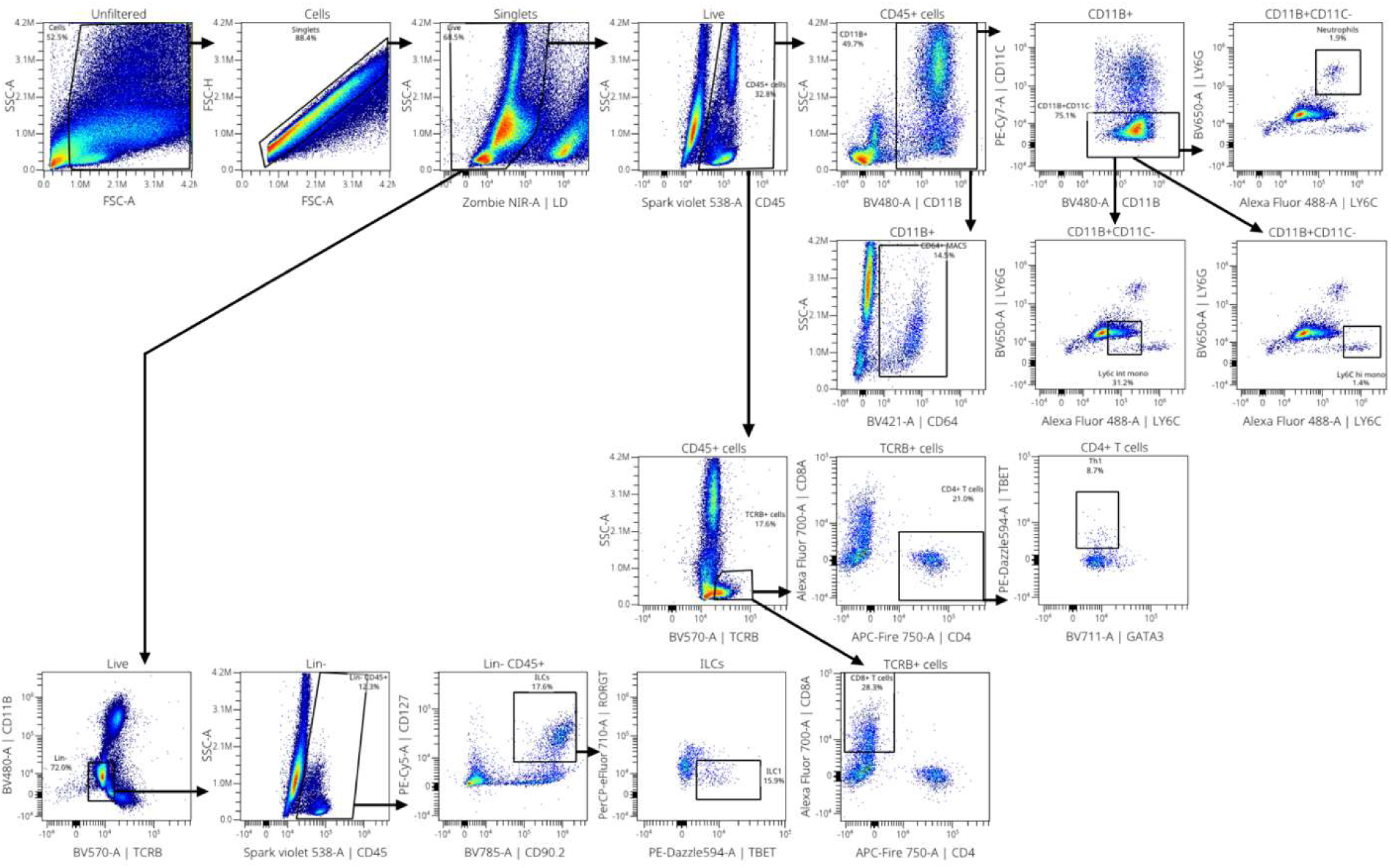
Flow cytometry gating strategy of the mouse colon. A representative flow cytometry gating strategy of a colon from a C57BL/6J mouse. Following SSC-A/FSC-A filtering and doublet exclusion pre-gating, Live CD45^+^ cells were split into TCRβ^+^ and CD11b^+^ gates to subsequently enumerate CD4^+^ T cells, CD8^+^ T cells, Ly6C^int^ monocytes, Ly6C^hi^ monocytes, and neutrophil populations. The CD4^+^ T cell pool was further divided according to TBET and GATA3 expression. To quantify ILC subpopulations, live CD45^+^ lineage-negative (Lin^-^; CD11b^-^ TCRβ^-^) cells were gated on CD127/CD90.2 and further segmented according to TBET and RORγT expression.

**Supplementary Fig. 2.**
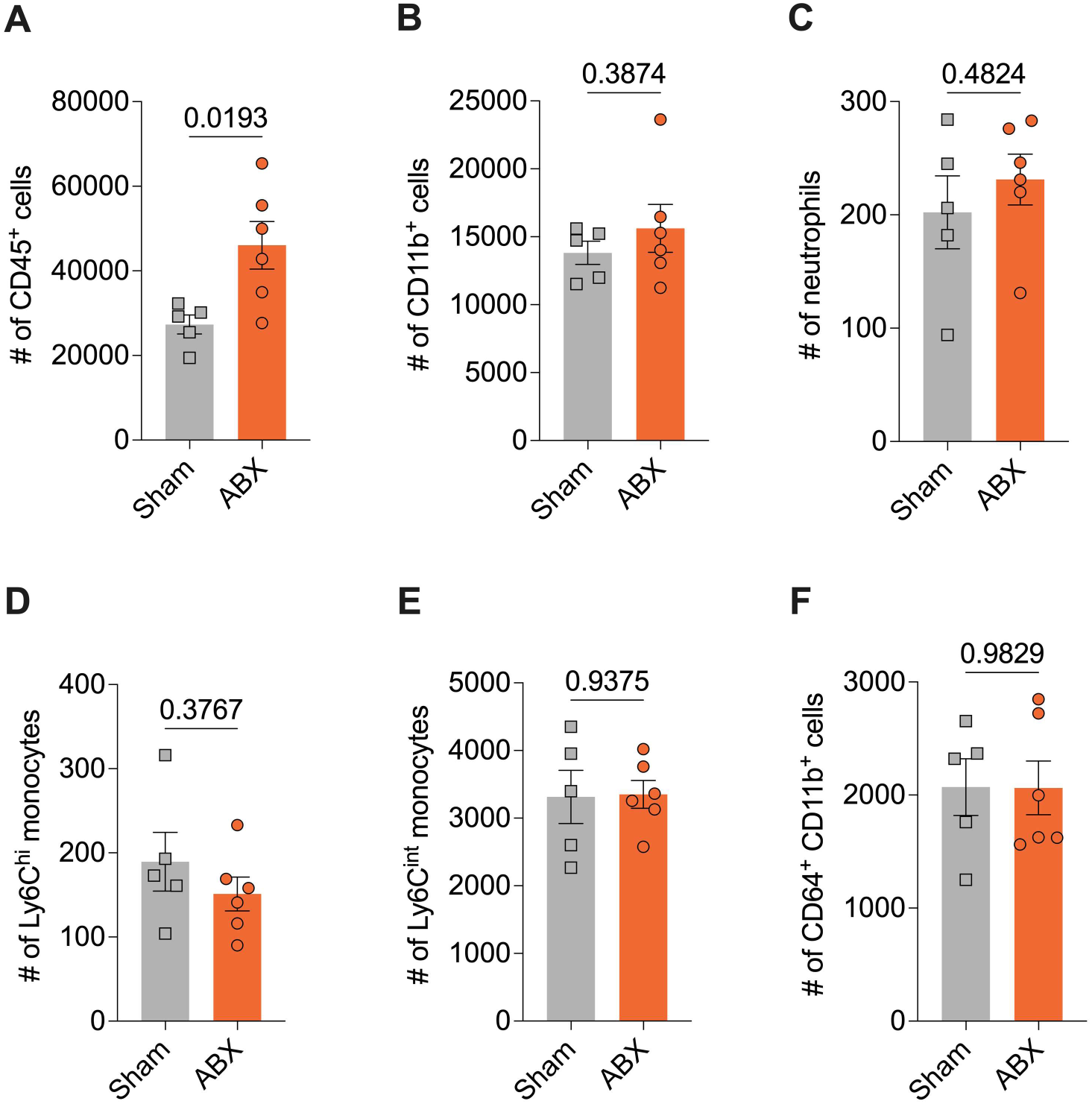
Colonic myeloid cell composition following ABX treatment. C57BL/6J mice received (1) either normal drinking water (sham treatment) or drinking water supplemented with an ABX cocktail of metronidazole, vancomycin, gentamicin, and colistin (ABX treatment) and (2) an i.p. injection of either a sham solution (sham treatment) or clindamycin (ABX treatment). The colon of each mouse was harvested 7 days after the start of the ABX regimen. (**A-F**) Cell count quantification of CD45^+^ cells (A), CD11b^+^ cells (B), neutrophils (C), Ly6C^hi^ monocytes (D), Ly6C^int^ monocytes (E), CD64^+^ CD11b^+^ cells (F) per harvested colon. Statistics calculated by a Welch’s t-test. Data are presented as mean (± SEM). Each symbol represents a value from an individual animal.

**Supplementary Fig. 3.**
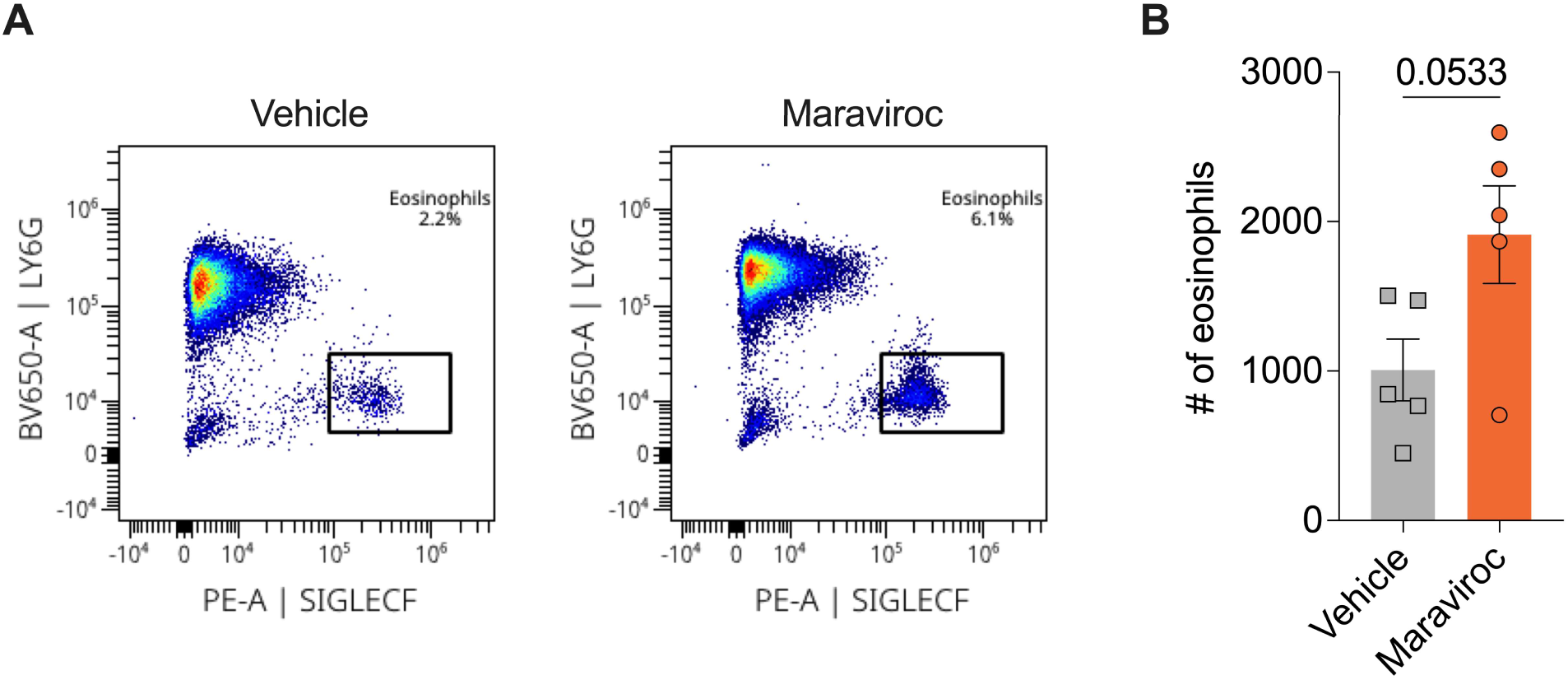
Therapeutic CCR5 inhibition promotes colonic eosinophilia during CDI. C57BL/6J mice received drinking water supplemented with an ABX cocktail of metronidazole, vancomycin, gentamicin, and colistin and an i.p. injection of clindamycin. Mice were infected with 1,000 *C. difficile* spores via oral gavage 24h after i.p. injection of clindamycin. On DPI-1, DPI 0, and DPI 1, mice were treated via i.p injections with either 1x PBS (vehicle) or a CCR5 inhibitor (maraviroc). The colon of each mouse was harvested on DPI 2 (7 days after the start of the ABX regimen). (**A**) Representative flow cytometry dot plots showing the gating strategy for eosinophils within the CD11c^-^ CD11b^+^ cell parent gate. (**B**) Cell count quantification of eosinophils per harvested colon. Statistics calculated by a Welch’s t-test. Data are presented as mean (± SEM). Each symbol represents a value from an individual animal.

## REFERENCES

1. Paredes-Sabja D, Shen A, Sorg JA. 2014. *Clostridium difficile* spore biology: sporulation, germination, and spore structural proteins. Trends in Microbiology 22:406–416.

2. Lessa FC, Mu Y, Bamberg WM, Beldavs ZG, Dumyati GK, Dunn JR, Farley MM, Holzbauer SM, Meek JI, Phipps EC, Wilson LE, Winston LG, Cohen JA, Limbago BM, Fridkin SK, Gerding DN, McDonald LC. 2015. Burden of Clostridium difficile Infection in the United States. New England Journal of Medicine 372:825–834.

3. Dubberke ER, Olsen MA. 2012. Burden of Clostridium difficile on the Healthcare System. Clin Infect Dis 55:S88–S92.

4. Guh AY, Mu Y, Winston LG, Johnston H, Olson D, Farley MM, Wilson LE, Holzbauer SM, Phipps EC, Dumyati GK, Beldavs ZG, Kainer MA, Karlsson M, Gerding DN, McDonald LC. 2020. Trends in U.S. Burden of Clostridioides difficile Infection and Outcomes. New England Journal of Medicine 382:1320–1330.

5. Kelly CP, Pothoulakis C, LaMont JT. 1994. Clostridium difficile Colitis. New England Journal of Medicine 330:257–262.

6. Bartlett JG, Chang TW, Gurwith M, Gorbach SL, Onderdonk AB. 1978. Antibiotic-Associated Pseudomembranous Colitis Due to Toxin-Producing Clostridia. New England Journal of Medicine 298:531–534.

7. Buffie CG, Pamer EG. 2013. Microbiota-mediated colonization resistance against intestinal pathogens. Nat Rev Immunol 13:790–801.

8. Deakin LJ, Clare S, Fagan RP, Dawson LF, Pickard DJ, West MR, Wren BW, Fairweather NF, Dougan G, Lawley TD. 2012. The Clostridium difficile spo0A Gene Is a Persistence and Transmission Factor. Infection and Immunity 80:2704–2711.

9. Lyras D, O’Connor JR, Howarth PM, Sambol SP, Carter GP, Phumoonna T, Poon R, Adams V, Vedantam G, Johnson S, Gerding DN, Rood JI. 2009. Toxin B is essential for virulence of Clostridium difficile. Nature 458:1176–1179.

10. Kuehne SA, Cartman ST, Heap JT, Kelly ML, Cockayne A, Minton NP. 2010. The role of toxin A and toxin B in Clostridium difficile infection. Nature 467:711–713.

11. Bishop EJ, Tiruvoipati R. 2022. Management of Clostridioides difficile infection in adults and challenges in clinical practice: review and comparison of current IDSA/SHEA, ESCMID and ASID guidelines. J Antimicrob Chemother 78:21–30.

12. Chen X, Katchar K, Goldsmith JD, Nanthakumar N, Cheknis A, Gerding DN, Kelly CP. 2008. A Mouse Model of *Clostridium difficile*–Associated Disease. Gastroenterology 135:1984–1992.

13. Reijnders TDY, Saris A, Schultz MJ, van der Poll T. 2020. Immunomodulation by macrolides: therapeutic potential for critical care. The Lancet Respiratory Medicine 8:619–630.

14. Franz T, Negele J, Bruno P, Böttcher M, Mitchell-Flack M, Reemts L, Krone A, Mougiakakos D, Müller AJ, Zautner AE, Kahlfuss S. 2022. Pleiotropic effects of antibiotics on T cell metabolism and T cell-mediated immunity. Front Microbiol 13.

15. Uddin MJ, Thompson B, Leslie JL, Fishman C, Sol-church K, Kumar P, Petri WA. 2024. Investigating the impact of antibiotic-induced dysbiosis on protection from Clostridium difficile colitis by mouse colonic innate lymphoid cells. mBio 15:e03338–23.

16. Okayasu I, Hatakeyama S, Yamada M, Ohkusa T, Inagaki Y, Nakaya R. 1990. A novel method in the induction of reliable experimental acute and chronic ulcerative colitis in mice. Gastroenterology 98:694–702.

17. Dorr P, Westby M, Dobbs S, Griffin P, Irvine B, Macartney M, Mori J, Rickett G, Smith-Burchnell C, Napier C, Webster R, Armour D, Price D, Stammen B, Wood A, Perros M. 2005. Maraviroc (UK-427,857), a Potent, Orally Bioavailable, and Selective Small-Molecule Inhibitor of Chemokine Receptor CCR5 with Broad-Spectrum Anti-Human Immunodeficiency Virus Type 1 Activity. Antimicrobial Agents and Chemotherapy 49:4721–4732.

18. Tan Q, Zhu Y, Li J, Chen Z, Han GW, Kufareva I, Li T, Ma L, Fenalti G, Li J, Zhang W, Xie X, Yang H, Jiang H, Cherezov V, Liu H, Stevens RC, Zhao Q, Wu B. 2013. Structure of the CCR5 Chemokine Receptor–HIV Entry Inhibitor Maraviroc Complex. Science 341:1387–1390.

19. Mencarelli A, Cipriani S, Francisci D, Santucci L, Baldelli F, Distrutti E, Fiorucci S. 2016. Highly specific blockade of CCR5 inhibits leukocyte trafficking and reduces mucosal inflammation in murine colitis. Sci Rep 6:30802.

20. Cowardin CA, Buonomo EL, Saleh MM, Wilson MG, Burgess SL, Kuehne SA, Schwan C, Eichhoff AM, Koch-Nolte F, Lyras D, Aktories K, Minton NP, Petri WA. 2016. The binary toxin CDT enhances Clostridium difficile virulence by suppressing protective colonic eosinophilia. Nat Microbiol 1:16108.

21. Buonomo EL, Cowardin CA, Wilson MG, Saleh MM, Pramoonjago P, Petri WA. 2016. Microbiota-Regulated IL-25 Increases Eosinophil Number to Provide Protection during Clostridium difficile Infection. Cell Reports 16:432–443.

22. Ganal SC, Sanos SL, Kallfass C, Oberle K, Johner C, Kirschning C, Lienenklaus S, Weiss S, Staeheli P, Aichele P, Diefenbach A. 2012. Priming of Natural Killer Cells by Nonmucosal Mononuclear Phagocytes Requires Instructive Signals from Commensal Microbiota. Immunity 37:171–186.

23. Gury-BenAri M, Thaiss CA, Serafini N, Winter DR, Giladi A, Lara-Astiaso D, Levy M, Salame TM, Weiner A, David E, Shapiro H, Dori-Bachash M, Pevsner-Fischer M, Lorenzo-Vivas E, Keren-Shaul H, Paul F, Harmelin A, Eberl G, Itzkovitz S, Tanay A, Santo JPD, Elinav E, Amit I. 2016. The Spectrum and Regulatory Landscape of Intestinal Innate Lymphoid Cells Are Shaped by the Microbiome. Cell 166:1231–1246.e13.

24. Zhang X, Borbet TC, Fallegger A, Wipperman MF, Blaser MJ, Müller A. 2021. An Antibiotic-Impacted Microbiota Compromises the Development of Colonic Regulatory T Cells and Predisposes to Dysregulated Immune Responses. mBio 12:10.1128/mbio.03335-20.

25. Ivanov II, Atarashi K, Manel N, Brodie EL, Shima T, Karaoz U, Wei D, Goldfarb KC, Santee CA, Lynch SV, Tanoue T, Imaoka A, Itoh K, Takeda K, Umesaki Y, Honda K, Littman DR. 2009. Induction of Intestinal Th17 Cells by Segmented Filamentous Bacteria. Cell 139:485–498.

26. Sefik E, Geva-Zatorsky N, Oh S, Konnikova L, Zemmour D, McGuire AM, Burzyn D, Ortiz-Lopez A, Lobera M, Yang J, Ghosh S, Earl A, Snapper SB, Jupp R, Kasper D, Mathis D, Benoist C. 2015. Individual intestinal symbionts induce a distinct population of RORγ+ regulatory T cells. Science 349:993–997.

27. Kim M, Galan C, Hill AA, Wu W-J, Fehlner-Peach H, Song HW, Schady D, Bettini ML, Simpson KW, Longman RS, Littman DR, Diehl GE. 2018. Critical Role for the Microbiota in CX3CR1+ Intestinal Mononuclear Phagocyte Regulation of Intestinal T Cell Responses. Immunity 49:151–163.e5.

28. Scott NA, Andrusaite A, Andersen P, Lawson M, Alcon-Giner C, Leclaire C, Caim S, Le Gall G, Shaw T, Connolly JPR, Roe AJ, Wessel H, Bravo-Blas A, Thomson CA, Kästele V, Wang P, Peterson DA, Bancroft A, Li X, Grencis R, Mowat AM, Hall LJ, Travis MA, Milling SWF, Mann ER. 2018. Antibiotics induce sustained dysregulation of intestinal T cell immunity by perturbing macrophage homeostasis. Science Translational Medicine 10:eaao4755.

29. Sugiyama K, Shirai R, Mukae H, Ishimoto H, Nagata T, Sakamoto N, Ishii H, Nakayama S, Yanagihara K, Mizuta Y, Kohno S. 2007. Differing effects of clarithromycin and azithromycin on cytokine production by murine dendritic cells. Clin Exp Immunol 147:540–546.

30. Iwamoto S, Kumamoto T, Azuma E, Hirayama M, Ito M, Amano K, Ido M, Komada Y. 2011. The effect of azithromycin on the maturation and function of murine bone marrow-derived dendritic cells. Clin Exp Immunol 166:385–392.

31. Lin S-J, Kuo M-L, Hsiao H-S, Lee P-T. 2016. Azithromycin modulates immune response of human monocyte-derived dendritic cells and CD4+ T cells. International Immunopharmacology 40:318–326.

32. Wilcox MH, Gerding DN, Poxton IR, Kelly C, Nathan R, Birch T, Cornely OA, Rahav G, Bouza E, Lee C, Jenkin G, Jensen W, Kim Y-S, Yoshida J, Gabryelski L, Pedley A, Eves K, Tipping R, Guris D, Kartsonis N, Dorr M-B. 2017. Bezlotoxumab for Prevention of Recurrent Clostridium difficile Infection. New England Journal of Medicine 376:305–317.

33. Peritore-Galve FC, Kroh HK, Shupe JA, Ehni AG, Cano Rodríguez R, Kordus SL, Washington MK, Knippel RJ, Stanley AM, Gamson A, Tkaczyk C, Lacy DB. 2025. The monoclonal antibody AZD5148 confers broad protection against TcdB-diverse Clostridioides difficile strains in mice. PLoS Pathog 21:e1013651.

34. Cowardin CA, Petri WA. 2014. Host recognition of *Clostridium difficile* and the innate immune response. Anaerobe 30:205–209.

